# Mathematical modeling of the frozen zone dynamics: towards using thermal imagers in cryotherapy

**DOI:** 10.1101/2024.10.19.619201

**Authors:** O. V. Ivakhnenko, O. F. Todrin, V.Yu. Globa, M.O. Chyzh, G.O. Kovalov, S.N. Shevchenko

## Abstract

Using thermal imaging for cryoablation and cryotherapy is a convenient technique but it does not allow for visualization of what is inside. We model thermal field distribution which can help relate surface thermal field images to in-depth temperature field dynamics that show freezing and thawing processes dynamics. We consider the spatio-temporal distribution of temperature using examples of hydrogel and living tissue subjected to a cryoprobe. Freezing in the hydrogel is compared with the measurements using visual morphometry. Such studies can be useful for comparing *in-vitro* and *in-vivo* thermal field dynamics and for estimating the correct timing for cryoapplication.

## Introduction

In various applications of cryotherapy, such as cryodestruction and cryostimulation, mathematical modeling of non-stationary temperature fields provides recommendations for the precise planning of the cryoablation time [1]. While cryotherapy remains largely empirical, modeling can predict and refine intuition by demonstrating different aspects of dynamics in living tissue, see [2–4] for review. Various theoretical aspects of modeling have been studied before such as freeze-thaw cryosurgical cycles [5], estimation of the stable frozen zone volume [6], thermal stresses appearing around a cryoprobe [7].

It is often more practical and ethical to use biological tissue gel-phantoms instead of experimenting on living tissue [8]. This approach allows for the utilization of the transparency of the hydrogel to observe the ice border position dynamics and compare it with numerical calculations. Also, hydrogel has consistent thermodynamic properties, so there is no need to collect huge statistics of the experiments.

Cryoapplication assumes movement of the freezing front, which is described by a Stefan-type free-boundary problem [3, 9, 10]. Mathematical modeling of thermal field dynamics in biological tissue must also take into account body heating to describe the metabolic heat generation and blood perfusion [11, 12]. As one important example of the latter, a thermally significant blood vessel presents a natural heat source, which competes with a heat sink (cryoprobe) [13, 14].

To perform cryosurgical procedures, one must precisely monitor the extent of freezing [2]. While local measurement techniques can be applied, such as inserting a thermal sensor, it usually disrupts the temperature field and leads to additional damage to the soft tissues. So the most used techniques include electrical impedance tomography, X-ray computed tomography, magnetic resonance imaging, and ultrasound imaging. The less used approach exploits thermal imagers, which provide an accessible and convenient tool [15, 16]. Thermal imagers can be used for monitoring even low temperatures [17], which makes them good for cryotherapy [18]. One of the problems that prevents the widespread use of thermal imagers for cryotherapy is that they provide only surface thermal-field images and do not give information about temperature distribution in bulk. We hope that this can be addressed by developing convenient modeling, to which we devote our present study.

So, “to improve cryosurgery further, there is the need to develop mathematical cryosurgery optimization techniques” [2]. To this end, we present mathematical modeling of freezing and thawing in both a biological tissue and a model medium gel-phantom. In doing so, we discuss modern calculation capabilities, such as using GPU, validate our calculations by comparing with measurements, and make conclusions on using thermal imaging for estimating the freezing volume and temperature field distribution inside the tissue or hydrogel.

Accordingly, the rest of the paper is organized as follows. In Sec. 1 we present basic equations for the free-boundary Stefan-like problem. Details of calculations, such as finite-difference method and heat flow continuity condition, are presented in Appendix A. Solutions for the hydrogel and living tissue are presented in Secs. 2 and 3 with the detailed parameters presented in Appendix B. Soving the heat equation with GPU is described in Appendix C. In Section 4 presents our experimental observations of the ice formation in the hydrogel. Then, in Sec. 5, we discuss using thermal images by showing how surface temperature image reflects three-dimensional freezing. In Sec. 5.3 we make conclusions and discuss further ways to improve this technique.

## 1 Formulation of the problem

To simulate the impact of the cryoapplicator on biological tissue or hydrogel we use the heat equation. For clarity, in what follows in this section, we will consider hydrogel, but the same model can equally be applied to biological tissues. In this work we consider the axial symmetry case to be able to eliminate the third dimension and consider only the radius and the depth in our temperature field simulation [11, 19, 20].

### 1.1 Heat equation

The thermal conductivity equation describes energy flow *q* through a material, which is proportional to the temperature gradient:

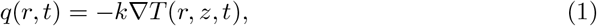

where *k* is the thermal conductivity, *T* is the temperature, and ∇ is the gradient of the scalar temperature field.

To describe heat distribution dynamics, we use the heat equation. This describes the temperature spreading in space over time:

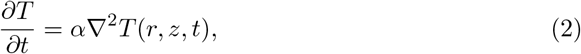

where

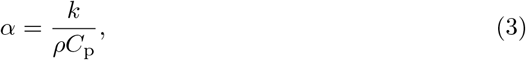

is the thermal diffusivity coefficient, *C*_p_ is the thermal capacity for constant pressure, *ρ* is the density, ∇^2^ is the Laplace operator. The Laplace operator in one-dimensional flat geometry is 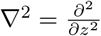. Then we consider the dependence of all thermodynamic parameters on temperature, which leads us to the non-uniform heat equation [21]

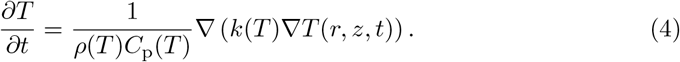

### 1.2 Stefan problem

Problems with dynamic phase change boundary usually are considered as a Stefan problem, Ref. [12]. Here changing the phase border position depends on the energy flow through the boundary. Energy flow is defined by the difference between the incoming and outgoing energy to the phase change border. For 1D flat geometry, it can be described by equation

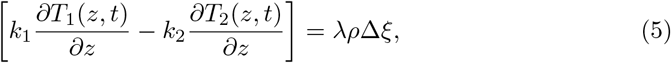

where *k*_1,2_ is the thermal conductivity coefficient of phases 1 and 2, *T*_1,2_ is the temperature of the respective phase, *r* is the coordinate, *λ* is the latent melting heat, *ρ* is the density of the material that changes phase, Δ*ξ* is the volume of the changing material at the boundary, it can be positive or negative due to energy flow in the current phase changing border, and leads to an increase or decrease in the phase change border position.

In our consideration we use a simplified approach to the Stefan problem by using the continuous heat equation, Eq. (4), including the latent phase change heat to the effective heat capacity, as described in Appendix B. This approach works well with solutions, where latent heat spreads on a wide range of temperatures and there is no exact phase transition temperature [22, 23].

### 1.3 Geometry and boundary conditions

Principal scheme of the cryo-application problem for 2D cylindrical geometry with axial symmetry, where the cryo-applicator with a temperature close to a boiling liquid nitrogen is pressed 1-2 mm into the hydrogel, is shown in Fig. 1. We assume that typical size of the frozen zone is significantly smaller than total size of the sample.

**Fig 1.**
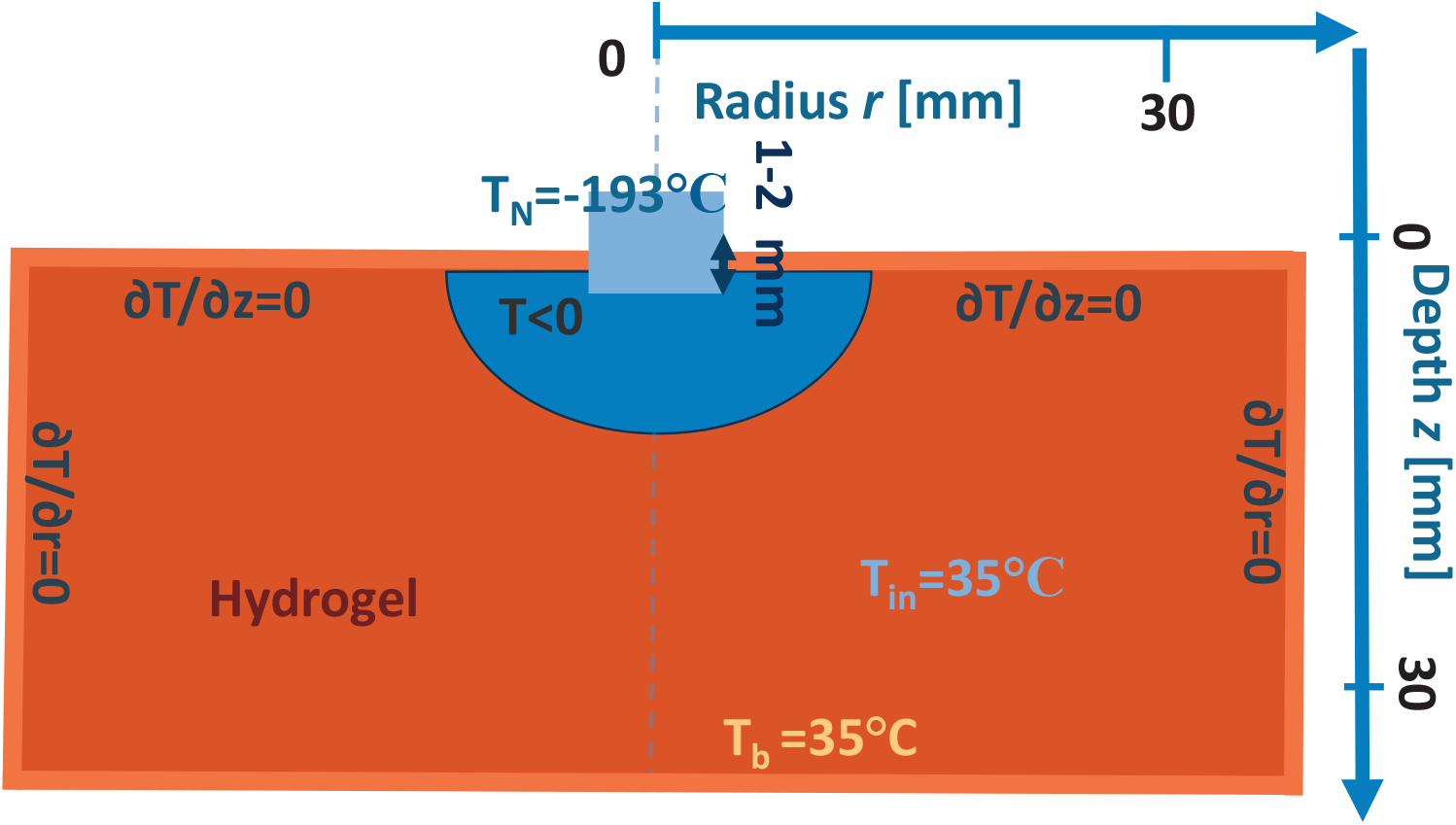
Scheme of the cylindrical geometry of the problem with freezing and thawing of hydrogel with initial and boundary conditions we used for the simulation.

1. Here we consider applying cryoapplicator with a tip radius *r*_ap_ = 4 mm. Which at coordinates less than *r < r*_ap_ has a constant temperature of the cryoapplicator *T*_N_ = −193^°^*C*.This condition is applied to all the time of freezing. Then during thawing stage we apply border condition at depth coordinate *z* = 0 heat flow is zero (no flow outside the system from zero depth coordinate).
  1.1 To have a more realistic case, we also consider pressing the cryoapplicator, which means the cryoapplicator is slightly pushed inside the hydrogel. That means we take the initial temperature in the area of radius of the cryoapplicator tip *r*_ap_ and depth *z*_ap_ = 1 mm and temperature *T*_N_ close liquid nitrogen boiling temperature. Also, we take into account a heat source far deep in the sample by assuming the heat bath at depth *z* = *z*_max_ with temperature *T*_b_ = 35^°^*C* degrees. The model also accounts for the heat flow from deeper tissue layers in the biological case. Here *z*_max_ is the maximum depth we take in numerical simulation.
  1.2 The cryoapplicator usually has no thermal insulation on its sides, which means that heat from the surface of the hydrogel also could dissipate through the air. That will lead to a bigger freezing circle on the skin or hydrogel than the simulation without air conductivity. However this additional freezing due to air conductivity is very shallow, so it disappears quickly after removing the cryoapplicator. For that, we take into account the thermal conductivity of air near the sample surface.
2. Boundary condition far deep from the considered freezing region temperature is constant *T*_b_ = 35^°^*C* degrees. The initial condition at phase boundary position *r > r*_ap_ is that the temperature is constant and equivalent to *T*_b_.
  2.1 Boundary conditions at the cryoapplicator border imply that there is a constant temperature *T*_N_, it assumes that cryoapplicator cooling productivity is significantly higher than heat flow to it.
3. After removing the cryoapplicator we replace the cryoapplicator volume with air and assume there is no heat transfer with air on the thawing stage. Freezing-stage border and initial conditions:
  - The cryoapplicator has a constant temperature *T*_N_ = −193^°^*C* close to the liquid nitrogen boiling temperature,
  - Initial temperature of the sample *T*_b_ = 35^°^*C*,
  - On the lower border temperature is kept constant at *T*_b_ = 35^°^*C*;

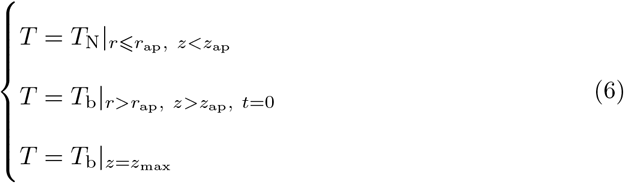 Thawing-stage border conditions:
  - No heat transfer through the sample-air interface,
  - Far deep inside the sample temperature is kept constant *T*_b_;

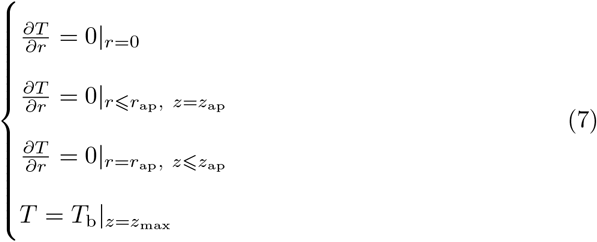

### 1.4 Numerical simulation

Heat dynamics problem with so large temperature difference and moving phase change border with complicated geometry, such as in this problem, is usually impossible to solve analytically [24]. So we use the finite difference method to solve the heat equation numerically [25].

The hydrogel and biological tissues do not consist of pure water but instead consist of solutions, which typically do not have a fixed freezing temperature. The solution usually freezes in some temperature range, for example from −0.1^°^*C* to −14^°^*C* for 5% gelatin hydrogel [8]. And phase change latent heat is distributed over this range of temperatures. Another feature of the hydrogel and the biological tissues is the significant temperature dependence of the thermodynamic parameters in so wide temperature range (from -193^°^C to 35^°^C), see Appendix B.

Taking into account both of these features we use the finite difference method to solve the heat equation with temperature-dependent coefficients throughout the simulation zone. We include in this model phase change latent heat as an additional heat capacity in the freezing temperature range. By using the finite difference method with temperature-dependent thermodynamic parameters, we also assume that the time step is small enough to keep thermodynamic parameters nearly constant during a single time step.

## 2 Frozen-zone dynamics in hydrogel

Hydrogel is an excellent model medium to investigate the impact of freezing on biological tissues because its thermodynamic parameters closely match those of biological tissues due to similar moisture concentrations. It allows us to obtain qualitatively similar results to those observed in biological tissues. However the main advantage is the simplicity of measuring thermodynamic parameters over a wide range of temperatures. So the thermodynamical parameters of the hydrogel are well investigated [8]. Another significant advantage of hydrogel is its consistent thermodynamic parameters, which allow to reproduce the impact of the cryoapplicator many times with high precision and adjusting theory and numerical calculations accordingly. In contrast to the hydrogel, thermodynamical parameters of the biological tissues can vary in a quite wide range and change from sample to sample due to different compositions by various factors such as hydration.

Using the parameters from Eq. (35) for the 5% gelatin hydrogel we perform simulation in the cylindrical geometry with Eq. (23), just using effective thermal capacity Eq. (38), Fig. 2. To solve the heat equation we use the finite difference method. Then we can compare our plots with ones, obtained by the thermal imaging technique in Refs. [16, 26]. Then we investigate what issue is caused by each of the typical stages they obtained. In Fig. 2 we show the result of our numerical calculations for hydrogel parameters for similar geometry as in Refs. [16, 26] with a 4 mm radius cryoapplicator pressed into the sample. In comparison to the experiment, we can also calculate temperature distribution in-depth and examine different isotherm dynamics, like -40^°^C isotherm, which temperature is considered lethal for the living cells, including cells of any malignant tumors.

**Fig 2.**
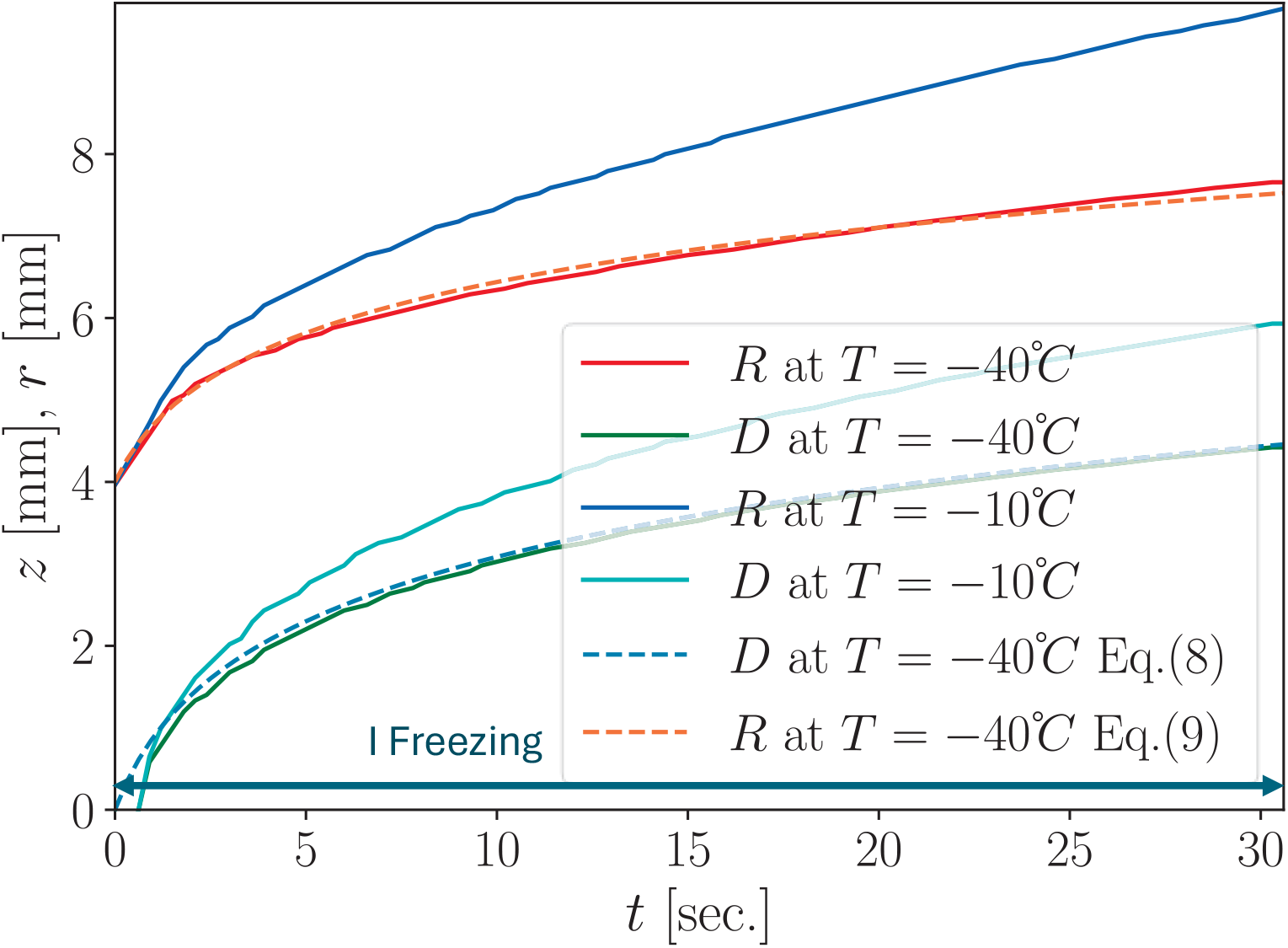
Numerical calculation of the maximum radius *R*(*t*) and maximum depth *D*(*t*) dynamics at the point *r* = 0 for two isotherms, − 0.1^°^*C* (which is the cryoscopic temperature of the hydrogel Eq. (31)) and − 40^°^*C* for 30 sec. of freezing time as in Refs. [16, 26]. We also show a comparison with the phenomenological formulas (dashed lines) for the depth and radius of the *T* = −40^°^*C* shown in formulas (8) and (9).

### 2.1 First stage: Freezing in hydrogel

During this stage cryoapplicator is applied. This leads to an ice spot size growth, which slows down over time. It happens due to the increasing surface of the contact with warmer hydrogel layers that lead to a decreasing of temperature gradient value, which corresponds to the cooling speed.

For the freezing of the hydrogel we found that the expanding of the ice spot radius as well as expanding isotherms like *T* = −40^°^*C* over time can be well approximated by natural logarithm. For example, depth for the *T* = −40^°^*C* in Fig. 2 can be well approximated as

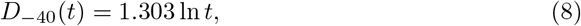

where *t* is the time in seconds and the value of *D*_−40_(*t*) is given in millimeters. For the radius of the *T* = −40^°^*C* isotherm we found

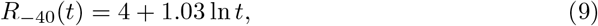

where 4 corresponds to the initial radius of the cryoapplicator in millimeters.

### 2.2 Thawing

During the thawing process, the cryoapplicator is removed and no longer provides a cold source, causing the ice spot to decrease in size due to heat flow from the thermal bath deep in a sample and surrounding material. Heat also transfers through the air, but this varies significantly with environmental conditions.

#### 2.2.1 Second stage: Fast decrease of frozen spot radius

The rapid decrease in the radius of the ice spot is caused by thermal conductivity, and thermal convection through the air, as well as the accumulation of frozen water vapor from the cold air around the cryoapplicator, during the freezing stage. To introduce it to the simulation we use effective parameters for the air, see Appendix A.5. This leads to the formation of a thin frozen circle on the sample surface surrounding the main ice spot, as shown in Fig. 4.

**Fig 3.**
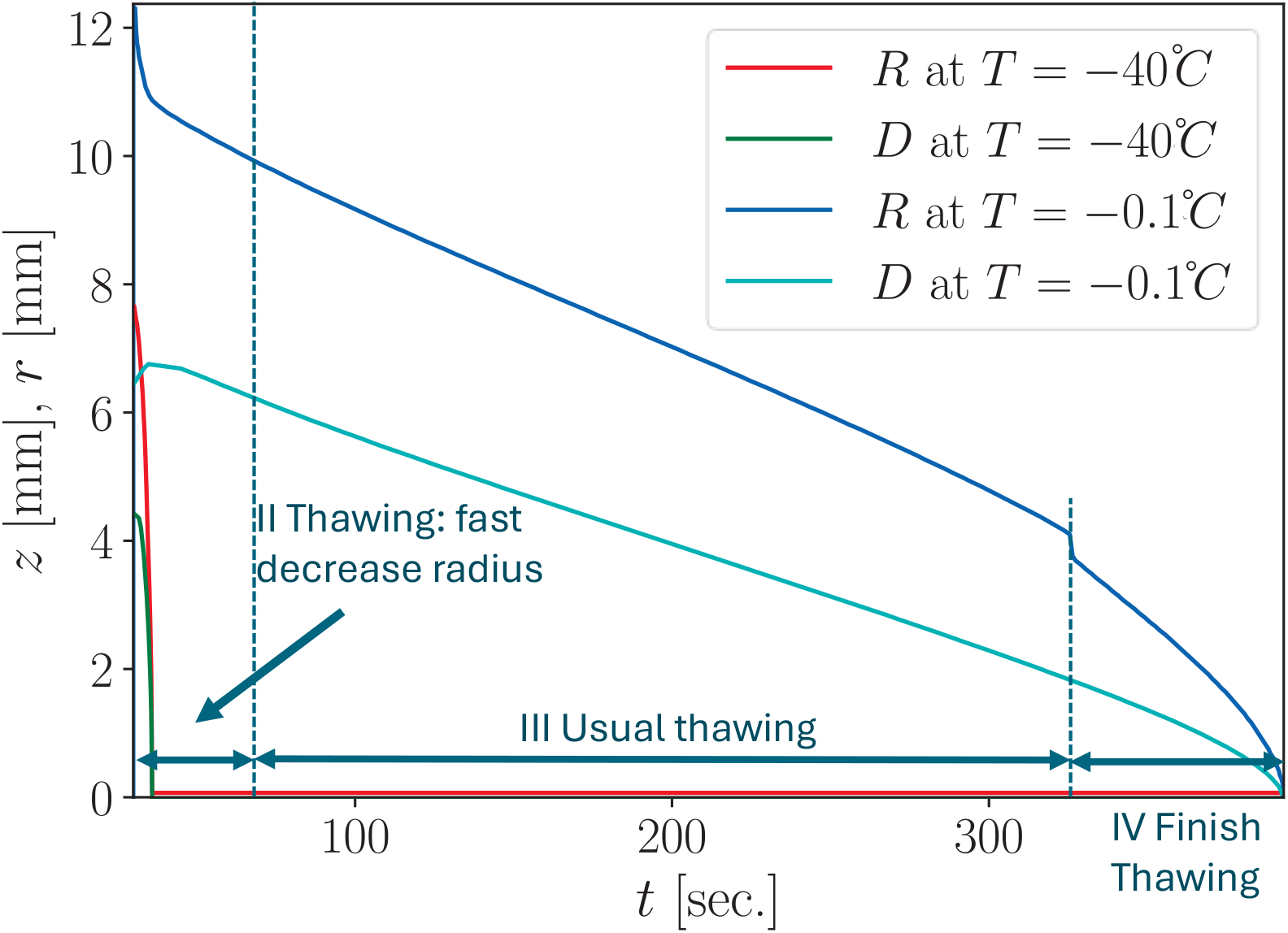
Numerical calculation of the maximum radius *R*(*t*) and maximum depth *D*(*t*) at the point *r* = 0 for two isotherms, − 0.1^°^C and − 40^°^*C*. For a 30 sec. of freezing time as in Refs. [16, 26], for the thawing stages.

**Fig 4.**
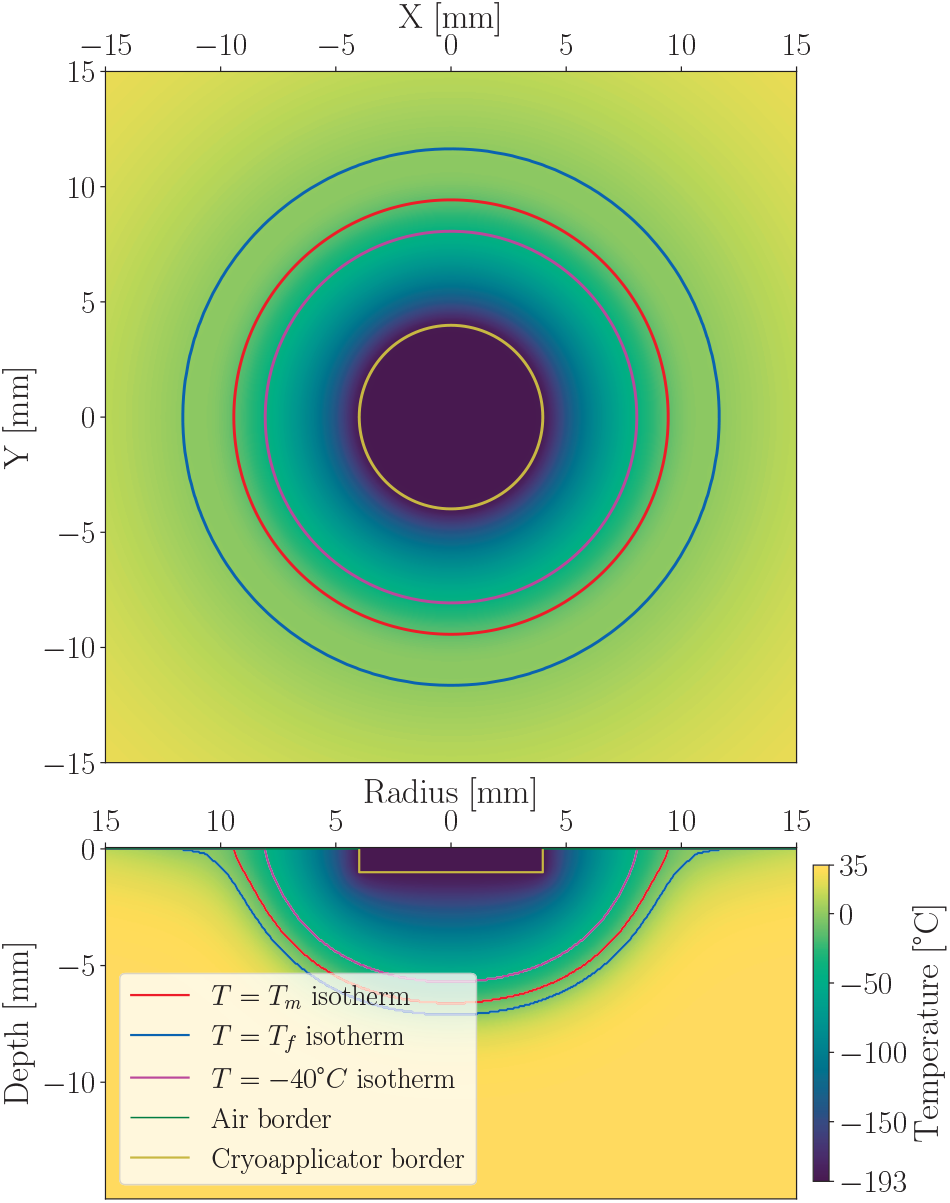
On the upper panel is the surface temperature field after 30 sec. of freezing. On the bottom panel is the temperature field distribution in depth. Several isotherm lines are shown: blue for *T* = *T*_f_ (upper limit of the phase change temperatures), red for *T* = *T*_m_ (lower limit of the phase change temperatures), and violet for *T* = − 40^°^*C* (death temperature for cells). The cryoapplicator border position is a yellow line. The frozen circle is formed around the main body of the ice spot. Here we include the heat transfer through the air, so you can see the thin layer of the frozen hydrogel around the main part of the ice spot on the zero temperature isotherm on the upper sides of the bottom panel. On the upper panel in cylindrical geometry with polar angle symmetry, the thermal image consists of circles with varying temperatures. By comparing the top and bottom panels we can see how the temperature field spreads in depth for the corresponding thermal image from the surface.

#### 2.2.2 Third stage: Main thawing

During this stage, the main ice spot is slowly thawing up to the radius of the cryoapplicator. Thawing is very slow, and the radius of the ice spot decreases gradually due to high latent melting heat. Additionally, the temperature inside the ice spot changes slowly due to latent heat being distributed over the temperature range between *T*_m_ and *T*_f_. Also, thermal conductivity of frozen tissues is significantly bigger than for thawed, it leads to very small temperature gradient inside the ice spot.

#### 2.2.3 Fourth stage: Final fast decreasing of radius

During this stage, the radius of the ice spot decreases rapidly due to the cryoapplicator was pressed during the freezing stage a few millimeters into the sample. The smaller the ice spot radius, the faster it decreases. It caused by the difference in volume between thawed and frozen hydrogel at the freezing front, this difference increases with decreasing the radius of the ice spot, i.e. 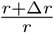 growth with *r* → 0. with the same heat flux due to a smaller radius and heat transfer surface respectively. At the beginning of this stage, the actual thickness of the ice spot under the cryoapplicator is thinner than the surrounding area due to the cryoapplicator was pressed a few millimeters inside the sample. As the radius of the frozen hydrogel reaches the radius of the cryoapplicator, the reduction in the ice spot’s radius accelerates by the thinner ice thickness under the cryoapplicator position.

## 3 Frozen-zone dynamics in biological tissue

Biological tissues are different from the hydrogel because they consist of many layers with different thermodynamic properties. During the application of the cryoapplicator to the skin, three main layers are involved: skin, underskin fat, and muscles, as shown in Fig. 5, as in Ref. [16].

**Fig 5.**
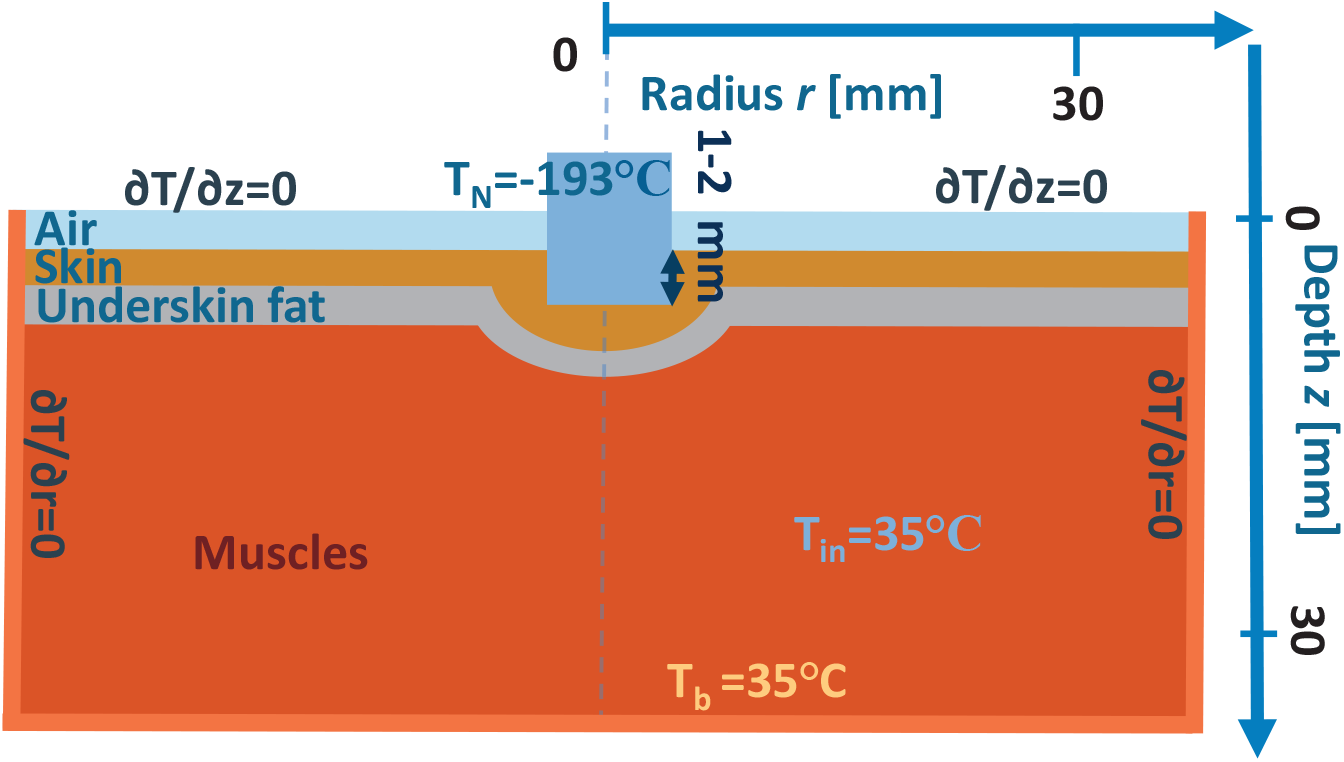
Material configuration for the biological tissues cryoapplication simulation, where orange is muscles, gray is under-skin fat, brown is skin, blue rectangular is cryoapplicator, and light blue is the air around the cryoapplicator.

### 3.1 Freezing

The frozen-zone dynamics are shown in Fig. 6. During freezing, due to the multi-layered structure of the biological tissues, we observe changes in the freezing speed, especially for the −40^°^C isotherm. Significant changes occur when tissues with markedly different thermodynamic parameters are included in consideration. Underskin fat could have a big impact on the spreading of the ice spot in depth slowing it down. It is caused by the significantly lower thermal conductivity of the underskin fat, especially when the temperature drops, and its thermal conductivity is several times lower than in other tissues as shown in Fig. 11. Increasing the thickness of the fat layer leads to decreasing the maximum reached the thickness of the ice spot as shown in Fig. 7.

**Fig 6.**
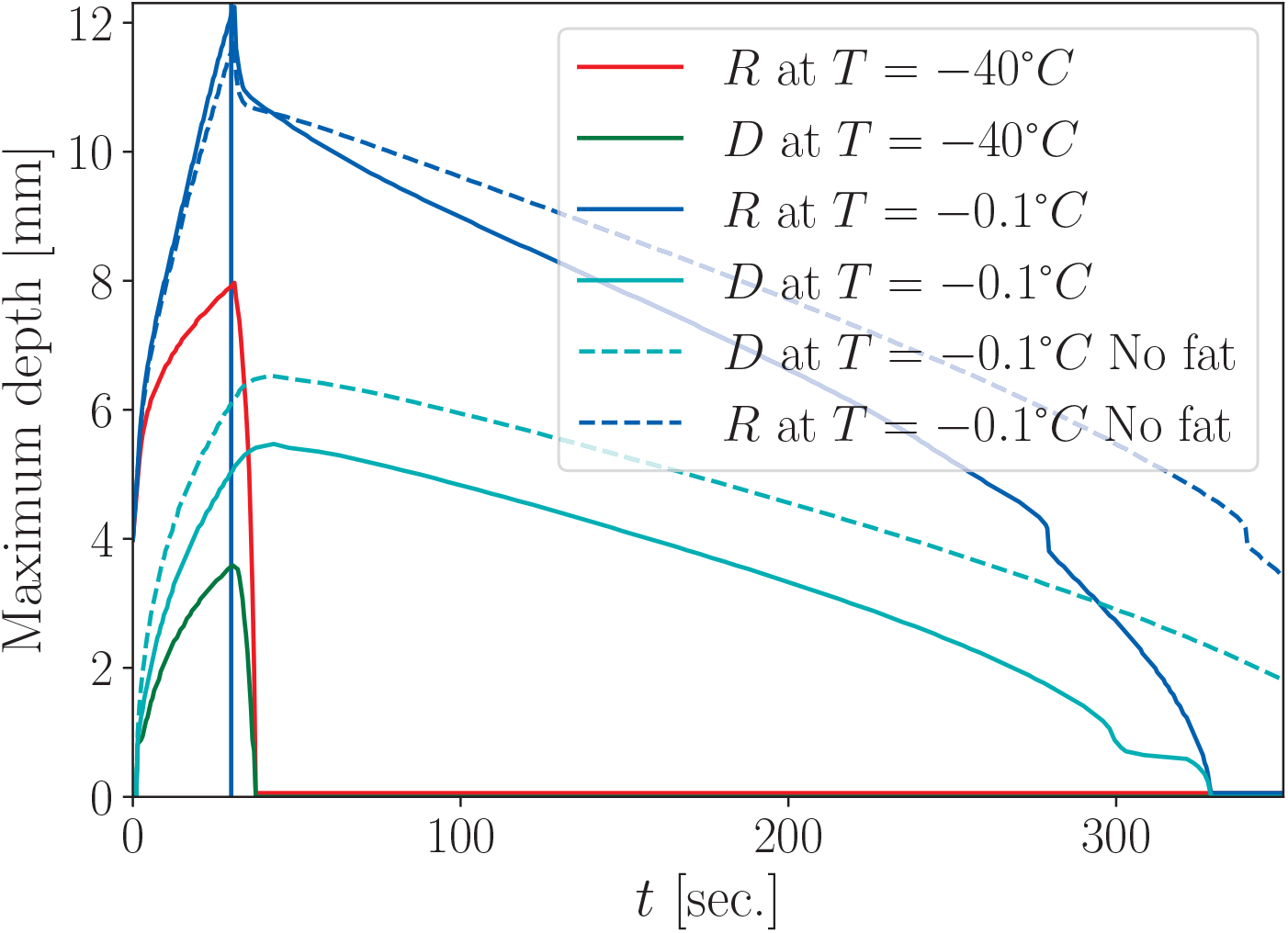
Numerical calculation of the maximum radius *R*(*t*) and maximum depth *D*(*t*) at the point *r* = 0 for biological tissues based on the three layers of the biological tissues, skin, underskin fat, and muscle layers is solid lines, dashed lines represent respective isotherms without underskin fat layer, for a 30 sec of freezing time as in Refs. [16, 26]. The dynamics for the maximum depth and maximum radius for two isotherms are shown for 0^°^*C* and − 40^°^*C*. The most noticeable difference here comes from the under-skin fat layer, which has significantly lower thermal conductivity, so the spread of the ice spot to the depth is slowed down significantly. Skin layer thickness is taken 0.6 mm, fat layer thickness is taken 0.3 mm for solid lines and 0 for dashed lines.

**Fig 7.**
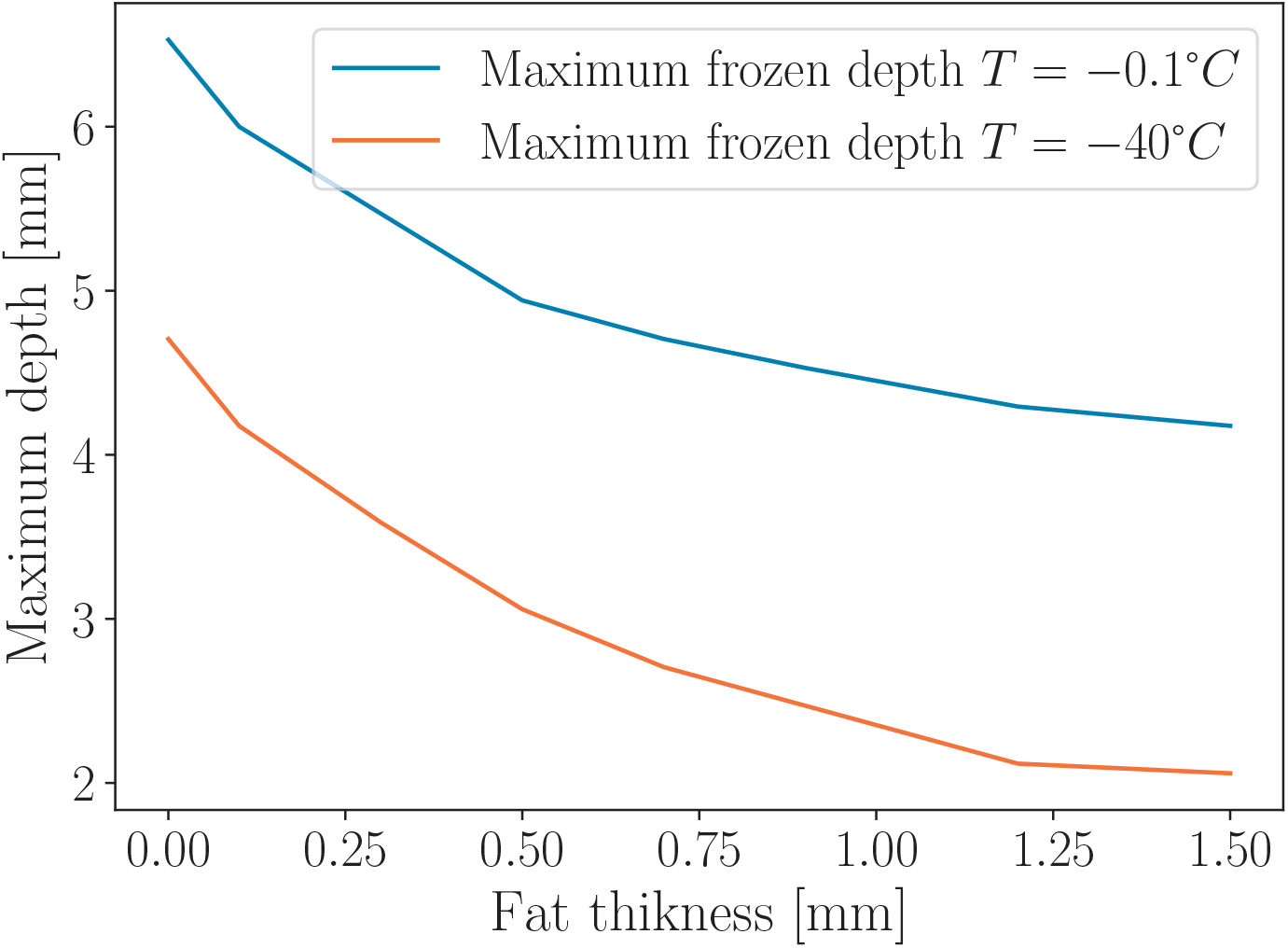
Numerical calculation of the maximum depth at the point *r* = 0 reached after 30 sec. of freezing for a different thickness of underskin fat in biological tissues based on the three-layer model with 0.6 mm skin layer, variable fat thickness, everything deeper is muscles. Here we show that with the growth in the thickness of the underskin fat layer maximum depth of the ice spot reduces due to the low thermal conductivity of the underskin fat. This describes the difference in the thawing times in Fig. 6 with the fat layer and without the fat layer, where a smaller maximum frozen depth leads to faster thawing. So applying a cryoapplicator aiming to destroy some tissues under a thick layer of fat or inside it can be difficult and need significantly more freezing time. It is caused due to significantly lower thermal conductivity of fat especially with lower temps as shown in Fig. 11.

### 3.2 Thawing in biological tissues

Thawing dynamics in the biological tissues can be similar qualitatively to the hydrogel when all the tissues have close thermodynamic properties, like skin and muscles. When we include the layer of underskin fat in our model, it begins to slow down the thawing process when the depth of the frozen spot reaches surface of the underskin fat at the end of thawing, as we show in Fig. 6 from 300 to 350 sec on the depth *D* at *T* = −0.1^°^*C* with the cyan solid line. Dashed line on the same figure for the zero underskin fat thickness almost does not have this horizontal segment. It is caused by that a thermal conductivity coefficient of fat is several times smaller than the one for other tissues, especially for lower temperatures, that is shown Fig. 11. Despite the fat slows down thawing, the overall thawing time for the case with fat is smaller than the one without fat, because the fat layer decreases frozen thickness, as shown in Fig. 7.

### 3.3 Surface thermal field imaging

Due to the limitations of the thermal imaging technique in the experiment, it is possible to see only the skin surface, but in our calculations, we can show the thermal image at any depth or angle inside the tissue that is shown in Fig. 4. So we plot the surface temperature thermal image in Fig. 4 to show comparable data to the thermal imaging experiment [16, 17].

## 4 Experimental frozen-zone dynamics in hydrogel

For verification of theoretical the calculations presented above, we did respective measurements. We studied the formation and dynamics of an ice hemiellipsoid *in vitro*. For this, we used a 5% gelatin hydrogel and the cryoapplicator with a radius of 4 mm. The radius and depth of the ice hemiellipsoid formed under the cryoapplicator were estimated from the data of visual morphometry. The full results of the experimental study of the dynamics of the movement of isotherms, which are important for cryotherapy (0, −20, −40), the shape and size of ice during prolonged low-temperature exposure with a flat applicator is only being prepared for publication; our main results on the formation of an ice hemiellipsoid are presented in Fig. 8.

**Fig 8.**
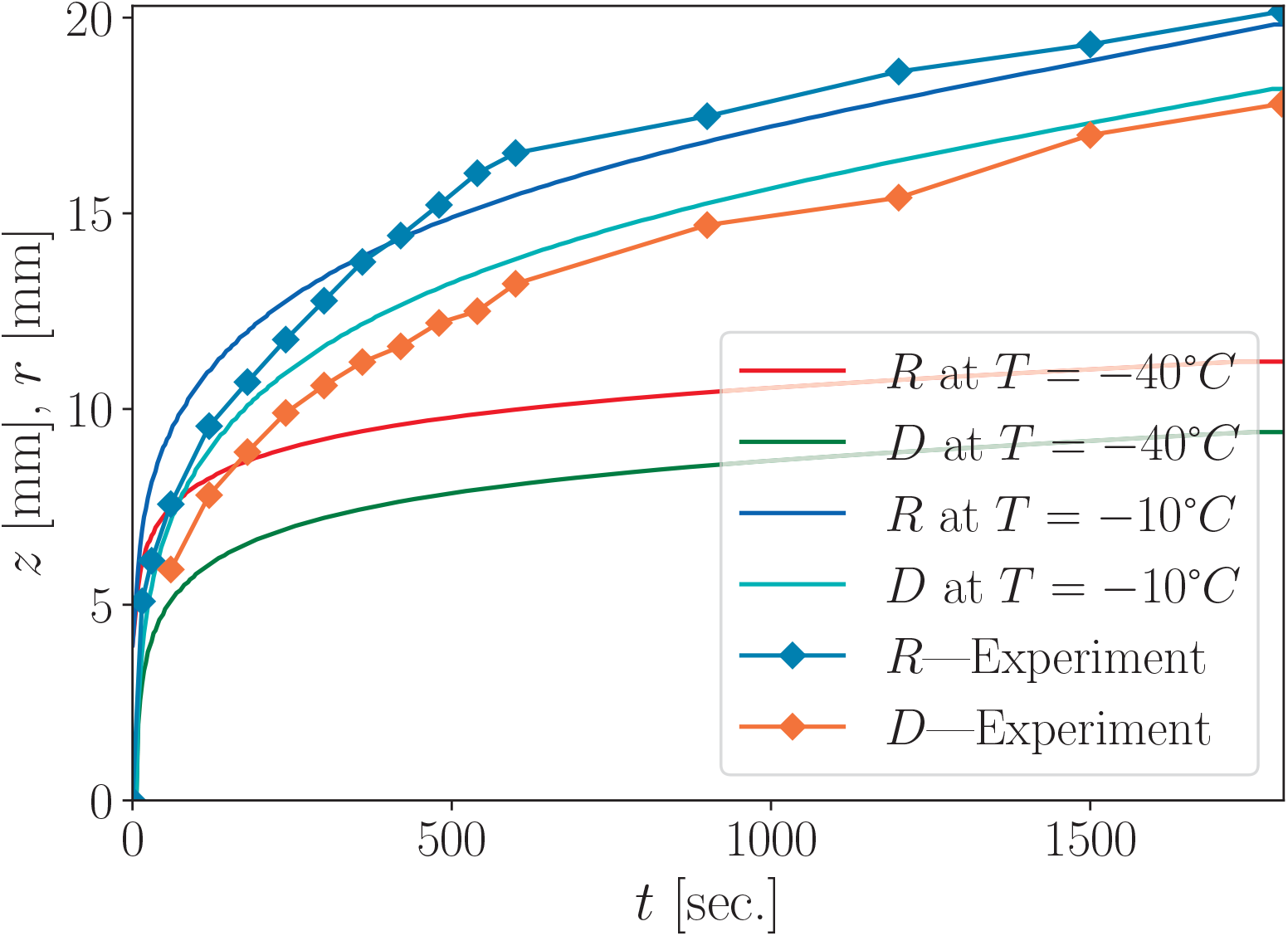
Formation of an ice hemiellipsoid in hydrogel under cryoapplicator. Experimental dependence of the radius and depth of the ice hemiellipsoid on time estimated from the data of visual morphometry is shown with square symbols. Respective numerical calculations, showing the radius and depth of the ice hemiellipsoid, which is the isotherm at *T*_*ice*_ = − 10^°^*C* that should correspond to the transparency change temperature in the hydrogel; these are similar to the data in Fig. 2 but for longer period of cryoapplication and for a different isotherms.

Dynamics of the radius and the depth of the ice hemiellipsoid, shown in Fig. 8, have a similar character with a rapid increase during the first ten minutes of the cryoapplication. To emphasize the dramatical increase, the radius was 6 mm after 15 seconds and was doubled in three minutes afterwards, while the increase during ten minutes starting from the 20*th* minute was only 3 mm. After half an hour the radius was 20 mm. The growth of the ice hemi ellipsoid in depth was somewhat delayed in relation to the radius. After half an hour the measured value of the maximal depth of ice was 17.8 mm. Thus, the relation of the depth to the radius was 0.78. So, we conclude that the cryoapplication to the 5% solution of the hydrogel gelatin results in the deformed ice hemi ellipsoid of the elliptic shape with the depth some twenty percent smaller than the radius on the surface.

Numerical calculation of the maximum radius *R*(*t*) and maximum depth *D*(*t*) at the point *r* = 0, in comparison with the experimental data for boundary with visual changes the transparency of the 5% gelatin hydrogel is shown in Fig. 8. Dynamics for the maximum depth and maximum radius is shown for two isotherms, −10^°^*C*, which is similar to the experimental data (the change of the transparency of the hydrogel happens at specific temperature inside the phase change region between *T*_m_ and *T*_f_), and −40^°^*C*. The difference between numerical calculations and the experiment can be caused by many factors, which were not taken into account in our simple heat transfer model, such as the formation of frost on the sides of the cryoapplicator, external heat sources, as well as model errors such as inaccuracies in the formulas describing thermodynamical parameters for the hydrogel, especially at cryogenic temperatures, the accuracy of the experimental measurements, etc. However, qualitatively they exhibit the same behavior mostly induced by the decreasing of the temperature gradient and lead to logarithm-like dependence of the radius and depth of the ice spot.

## 5 Discussion of application for cryoablation

A mathematical model can be useful for predicting the size of certain isotherms in an ideally controlled environment for biological tissues. However, in experiments, many factors can influence thermal dynamics, leading to some differences, although the general qualitative trend of dynamics should remain similar.

### 5.1 Is gel good for describing freezing/warming of living tissue?

The gel is a good model medium for simulating the cryoapplication problem when compared with mostly uniform biological tissues [8], but it demonstrates differences in behavior when layers of biological tissues with significantly different thermal properties. Another advantage of using hydrogel is its well-investigated and consistent thermal properties, unlike living biological tissues, which have varying thermal properties due to various factors.

### 5.2 How can surface termograms make conclusions about what’s inside?

Thermal imaging is a very useful tool for calibrating numerical calculations by providing reference points on the surface temperature field. This was demonstrated in Figs. 4 and 8. In particular, we concluded that the frozen zone has an elliptic shape with the surface radius (visualized by the thermal imager) around 20 percent larger than the depth (which is the value targeted in cryoablation).

### 5.3 Practical hints on using thermal imaging for cryoablation and cryotherapy

By knowing the thermal properties and thickness of the different layers and reducing temperature leakage through the air, thermal imaging can be used to evaluate the depth of certain isotherms. Using our numerical approach. The structure of the biological tissue can significantly influence the thermal distribution and dynamics as we demonstrated in Fig. 6.

## Conclusions

We presented a practical and self-sufficient description of how to mathematically model thermal distribution for problems involving the use of a cryoapplicator. Using this model, we studied freezing and thawing in two different media—hydrogel and living tissue—under realistic parameters. Our calculations of the spatio-temporal thermal field distribution were used to describe experiments on a gelatin gel [26] and on the living bodies of rats [16]. Validating our calculations with the measurements allows us to draw the following conclusions.

1. *In vitro* measurements on 5% gelatin hydrogel can be used not only for qualitative but also for quantitative comparisons with *in vivo* measurements. Simulation on a hydrogel phantom of freezing and melting during cryosurgery is a good alternative to animal experiments.
2. When modeling freezing and thawing in gels, we found that the freezing zone has a hemiellipsoid shape with maximal depth some 20% smaller than the surface radius for the 4 mm radius cryoapplicator.
3. When modeling freezing and thawing in living tissue, we found that the significant thermodynamic parameters difference between layers in the multi-layered structure (e. g. the underskin fat layer with low thermal conductivity) can cause deviations from the homogeneous situation, like in the hydrogel.
4. When interpreting the results of modeling freezing and thawing processes in hydrogels, it is necessary to take into account the features of the thermophysical parameters of a specific zone of a biological object in which cryoapplication will be performed. These features, in particular, are determined by the thickness and presence of layers of various biological tissues, as well as heat influxes from blood vessels, which depend on the proximity of the vessels to the cooling zone and their number, the diameter of the vessels and the blood flow velocity in the vessels at different times of cryotherapy.
5. Our results show how to correlate internal thermal dynamics with a surface thermal image. Using the correspondence between surface and internal thermal dynamics in the cryosurgery zone makes it possible to predict the shape and size of the freezing zone and the distribution of isotherms that are important for cryosurgery in the depth of tissues directly during cryoapplication. Even though the thermal imager records the temperature distribution only on the surface, continuous visualization of thermal fields during cryoapplication makes it possible to evaluate the dynamics of isotherms (−20^°^*C* and −40^°^*C*) that are critical for guaranteed destruction of biological tissues. Monitoring the dynamics of different isotherms opens up the possibility of timely changing the parameters of low-temperature exposure in such a way as to minimize the undesirable consequences of excessive cooling of biological tissues, provided that guaranteed cell death in the pathological tissue is achieved.

## Acknowledgments

The authors acknowledge the useful discussions with G.V. Shustakova, Y.V. Fomenko, E.Y. Gordienko, and the technical assistance of M. Liul, P. Kofman, A. Ryzhov. This work was supported by the National Research Foundation of Ukraine (Grant No. 2022.01/0094).

## A Method of finite difference

The Finite Difference Method (FDM) is a numerical technique used to approximate solutions to differential equations [25], such as the heat equation. The main disadvantage of this method is its potential stability issues, particularly when dealing with nonlinear derivatives. To address this, smaller time and spatial steps are often required. FDM solves the heat equation on the mesh of points, so the main parameter of this method is a number of points, which is determined by the size of the problem divided by the spatial step Δ*x* and time step Δ*t*.

### A.1 Flat geometry

Consider the mesh with flat geometry. In this mesh, the approximate linearized derivative at each node can be calculated as the right and left derivatives, denoted as *T* ^+^, *T* ^−^, respectively,

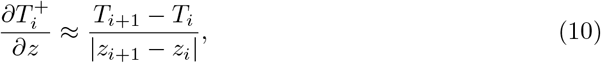

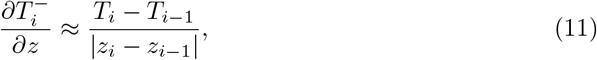

where *z* is the flat geometry coordinate. Then consider the heat equation (1) and calculate the total heat changing in one node on the mesh

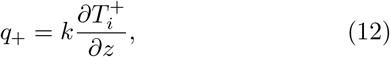

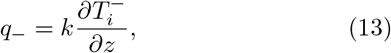

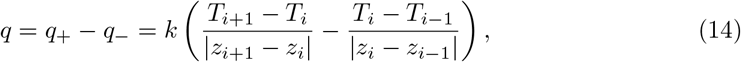

where *k* is the thermal conductivity coefficient; in a realistic case, it also significantly depends on the temperature. Here 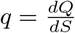 is the density of the heat flux, *q*_+_ and *q*_−_ is the heat flux to each direction so *q* = *q*_+_ + *q*_−_. Where *Q* is the heat energy, *S* is the surface area that heat energy passes through. In the case of equidistant flat geometry |*z*_*i*+1_ − *z*_*i*_| = |*z*_*i*_ − *z*_*i*−1_| = Δ*z*, mesh for the time Δ*t* = *t*_*i*+1_ − *t*_*i*_, we obtain changing of the heat flux:

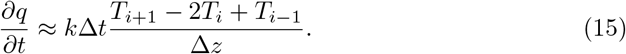

This equation becomes a true equation only with infinitely small value of time step *dt*. Therefore for simplicity, we consider heat spreading inside each phase far from the phase-changing border. It means that heat, obtained by the element of the mesh leads to only changing the temperature of that node with volume *dV* = *dSdz*. So we obtain:

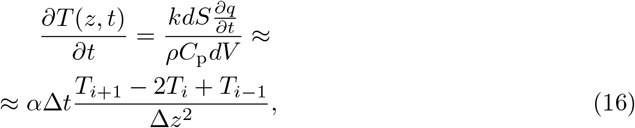

where 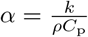 is the thermal diffusivity coefficient.

### A.2 Radial geometry

In radial geometry, the derivative is defined differently, with the gradient operator given by 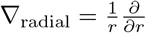. To calculate the derivative we use a half-shifted radius to precisely calculate the derivatives between midpoints, which is important in the radial geometry,

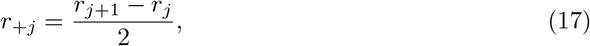

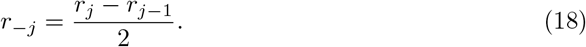

So the heat flow in radial geometry

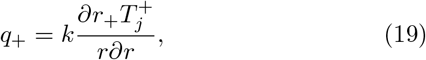

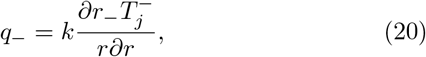

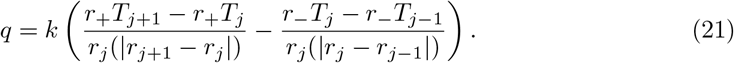

As a result, we obtain heat equation in radial coordinate

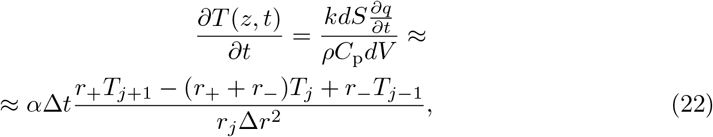

where Δ*r* = *r*_*j*_ − *r*_*j*−1_ is the radius mesh step.

### A.3 Cylindrical geometry

Cylindrical geometry can be modeled by combining the flat geometry for the depth coordinate *z* and the radial geometry for the radial coordinate *r*. This approach results in the following heat equation in cylindrical coordinates with polar angle symmetry (i.e. no dependence on the polar angle):

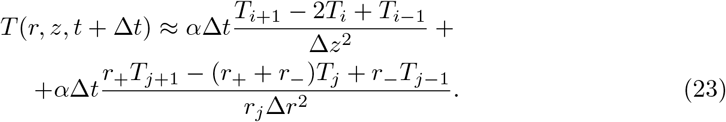

### A.4 Heat flow transfer for *α*(*T*)

When the thermal diffusivity coefficient *α* depends on temperature, it is crucial to ensure that the heat flow remains continuous, assuming no heat sources or sinks are present at specific nodes. It means that the amount of heat transferred to a neighbor node should be equivalent to the heat the neighbor node receives. For this, the mutual conductivity between every couple of neighbor nodes should be the same, which means that we cannot use just heat diffusivity in each node, we should differentiate heat diffusivity between each pair of nodes and calculate the effective thermal diffusivity coefficient in each direction of heat transfer

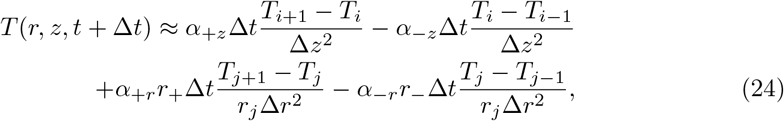

where

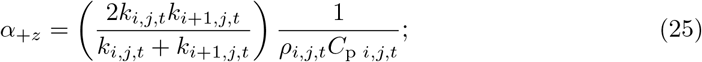

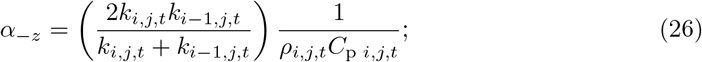

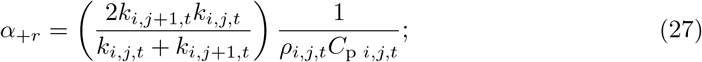

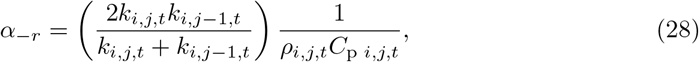

*k*_*i,j,t*_ is the thermal conductivity in the *i, j* node number in moment of time *t*. Here, *i* corresponds to the node number along the *z* coordinate, *j* corresponds to the node number along the radial coordinate *r*, and *t* represents the time step number. We use the analogy of parallel resistors to compute the effective thermal conductivity, given by

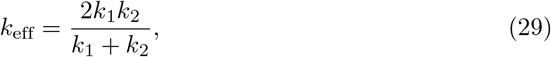

which is equivalent to consecutive thermal conductivities for two layers with 0.5*ξ* thickness. Here we assume that half the distance between nodes corresponds to one node’s material and temperature, second half corresponds to the neighbor node. Here *ξ* refers to the distance between nodes *δr* for radial coordinate or by *δz* for depth. As you can notice from the Eq. (29) the distance between the centers of each node refers to *ξ* and within this distance we have that half of this distance corresponds to one node, and the second half corresponds to the neighbor node. That is shown in Fig. 9.

**Fig 9.**
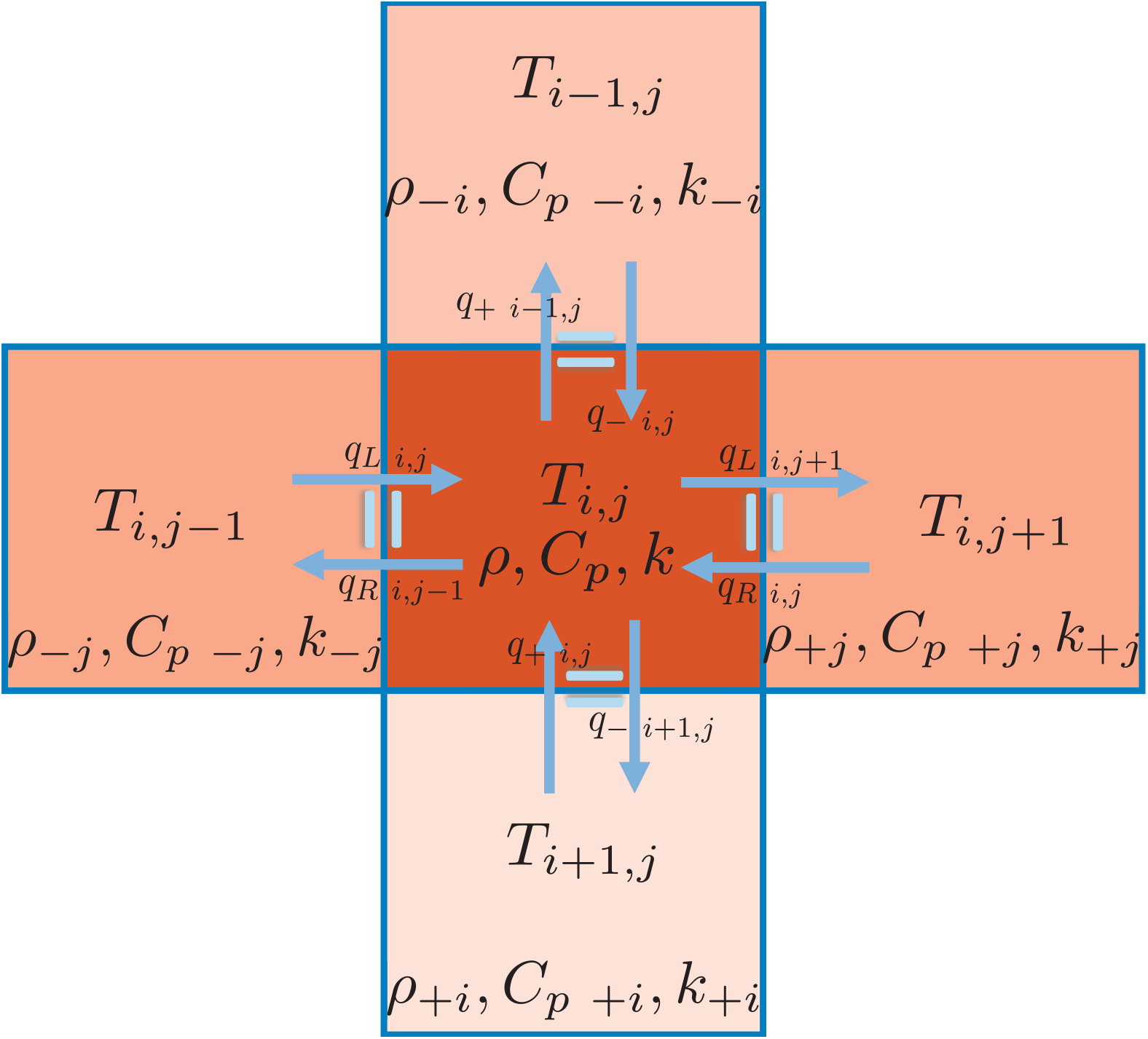
Scheme of nodes in FDM, where we show the heat transfer equality in all directions to the four neighbors. There are different temperatures in the different nodes, as well as different temperature-dependent parameters: density *ρ*, thermal conductivity *k*, and effective heat capacity *C*_p_; so we consider neighbor nodes as different materials and calculate respective thermal conductivity for the multilayer structure in each direction by Eq. (29).

### A.5 Heat transfer through the air

When we consider thermal conductivity through the air we try to take into account several processes such as air thermal conductivity, convective heat exchange between the cryoapplicator with non-insulated sides and skin, condensation of water vapor from the air on the cryoapplicator sides and skin. To simulate all these complicated phenomena and include them in the FDM simulation we need to make approximations. If we just use thermodynamical parameters of the air, without simulating heat convection and frost forming, the air level above the skin will just reach the same temperature as the skin very fast and will not provide additional cooling, because the density of the air is a thousand times less than hydrogel or biological tissues density. For this, we use the effective parameters to describe all phenomena of heat transfer through the air. We use adjustable thermal conductivity inside the air layer, different thermal conductivity in the direction of the skin, and adjustable heat capacity. Another important parameter is the thickness of the air layer, which also influences the heat transfer in our model. Next, we perform simulations to find and adjust effective parameters by comparison with the cryoapplication experiment on rats [16, 26], from which we observe the additional radius of the thin layer around the main ice spot, and compare it with that we obtained in our calculations.

## B Parameters

### B.1 Parameters of the hydrogel

As a model material, we use 5% gelatin gel, with parameters taken from Ref. [8]:

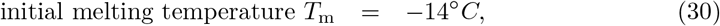

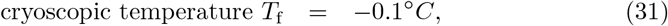

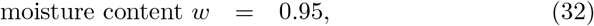

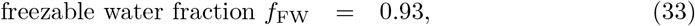

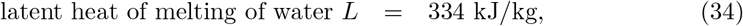

the temperature-dependent density *ρ*

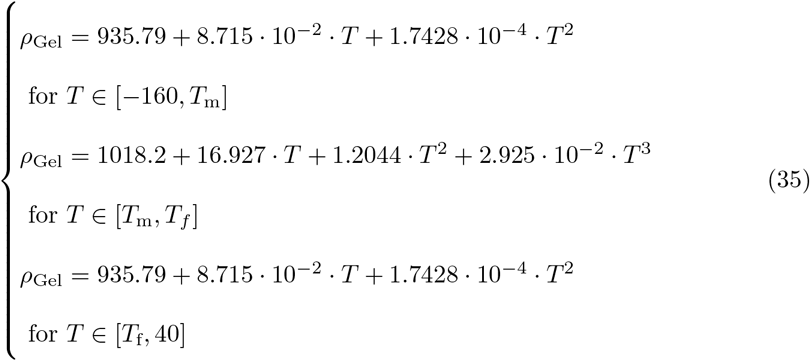

temperature-dependent capacity *C*_p_ and temperature-dependent thermal conductivity *k*:

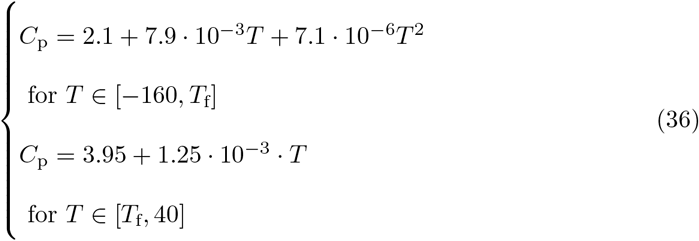

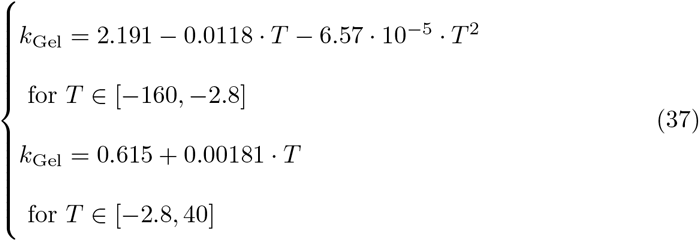

A key feature of solutions freezing is the change in water concentration in the unfrozen part because ice preferentially forms from pure water, leading to a decrease in the freezing temperature of the liquid part of the hydrogel or biological tissues.

To account for the latent heat of phase change, we introduce effective thermal capacity [27] in the temperature range between *T*_m_ and *T*_f_ :

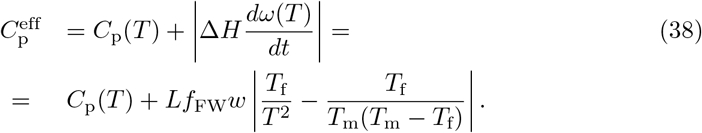

### B.2 Parameters of the biological tissues

Here, we present the parameters for biological tissues. All thermodynamic parameters of biological tissues depend on temperature, but there is significantly less data due to the variability and inconsistency of parameter values between different samples. This is due to the wide range of factors affecting biological tissues, one of the most significant is hydration, which corresponds to the moisture content level in the tissues.

From Table 1 we can see that the temperature range of the phase transitions for the skin and muscles is very similar. The parameters of the under-skin fat are significantly different due to its lower moisture level.

**Table 1.** Thermodynamic parameters for biological tissues. Since the thermodynamic parameters for the same type of tissues are similar among mammals, [28–30].

| Material | $L$ [kJ/kg] | $T_f$ [°C] | $T_m$ [°C] |
| --- | --- | --- | --- |
| Skin | 230 | −0.53 | −9.15 |
| Skin Fat | 100 | −5 | −10 |
| Muscle | 250 | −0.91 | −10 |

In Table 2 and Fig. 10 we can see that the density of all biological tissues is quite similar across a wide temperature range.

**Table 2.** Density of the biological tissues *ρ* [kg/m^3^], [31, 32].

| Material | $T > T_f$ | $T < T_f$ |
| --- | --- | --- |
| Skin | 1100 | 1039 |
| Skin Fat | $986.47 - 0.44503T$ | 978.6 |
| Muscle | 1047 | 988.3 |

**Table 3.**
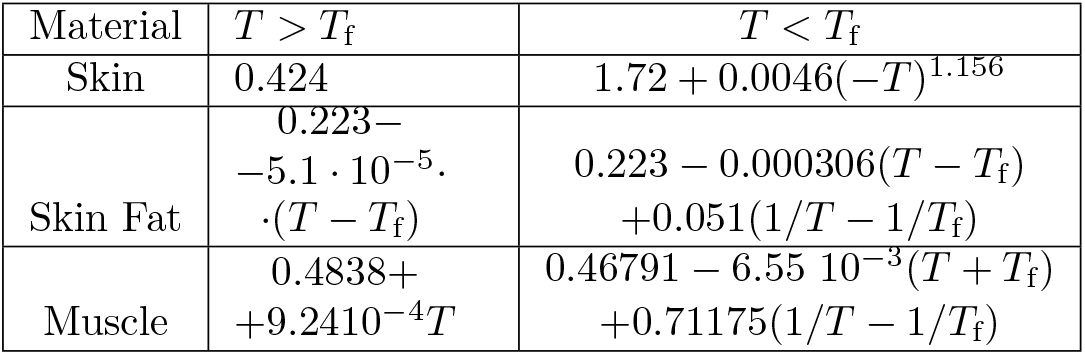
Thermal conductivity for the biological tissues *k* [Wt/(m K)], [31, 33, 34].

| Material | $T > T_f$ | $T < T_f$ |
| --- | --- | --- |
| Skin | 0.424 | $1.72 + 0.0046(-T)^{1.156}$ |
| Skin Fat | $0.223 - 5.1 \cdot 10^{-5} \cdot (T - T_f)$ | $0.223 - 0.000306(T - T_f) + 0.051(1/T - 1/T_f)$ |
| Muscle | $0.4838 + 9.2410 \cdot 10^{-4}T$ | $0.46791 - 6.55 \cdot 10^{-3}(T + T_f) + 0.71175(1/T - 1/T_f)$ |

**Fig 10.**
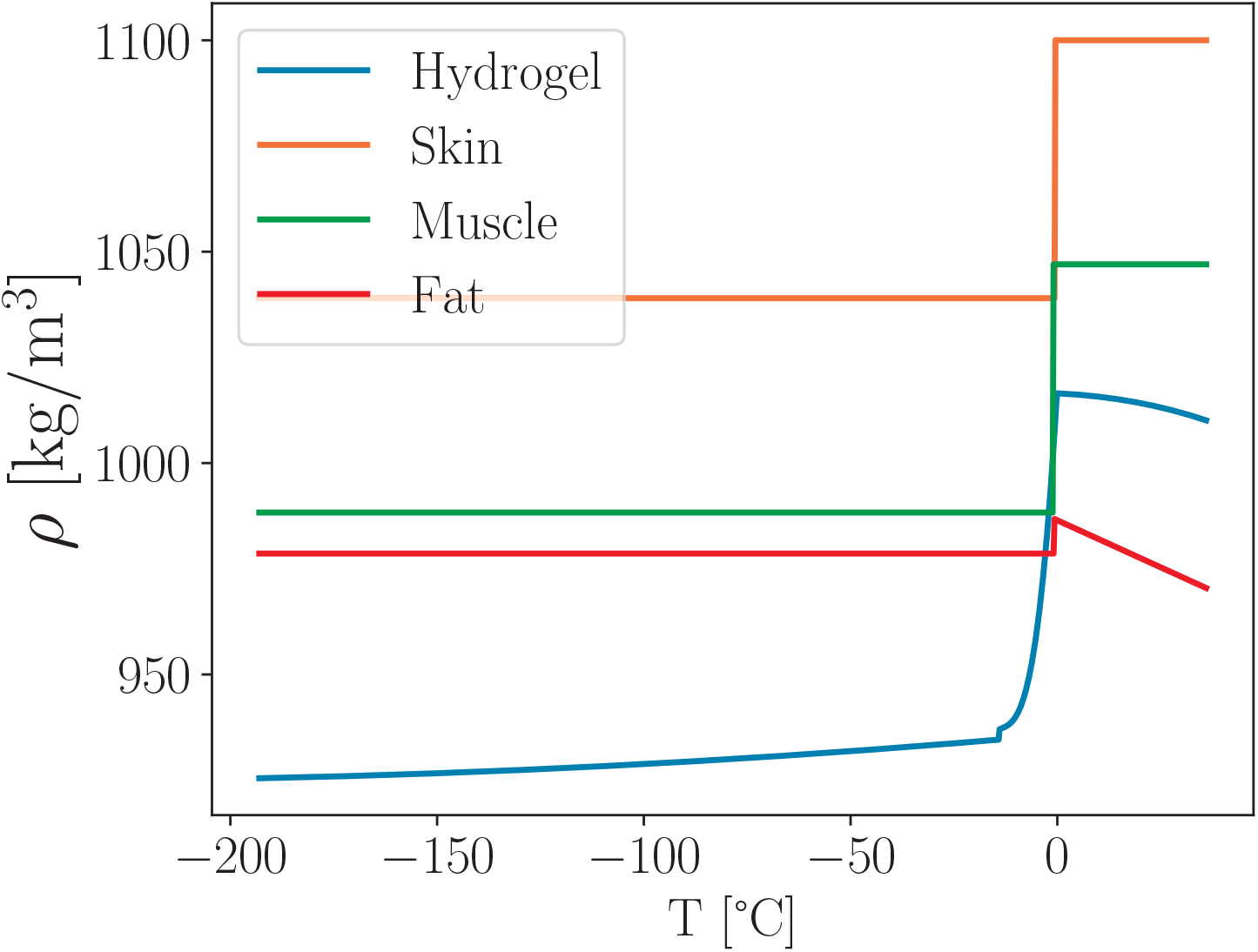
Temperature dependence of the density of the biological tissues and hydrogel, plotted based on Table 2 and Eq. (35).

In Table 3, we can see that the thermal conductivity of the biological tissues is defined by different formulas, it is because these data were collected from several different sources, that use different polynomial approximations to fit the experimental data [31, 33].

In Fig. 11 we can see that the thermal conductivity of underskin fat is significantly different from that of skin and muscles. This difference leads to significant changes in the temperature field when underskin fat is included in the calculation model.

**Fig 11.**
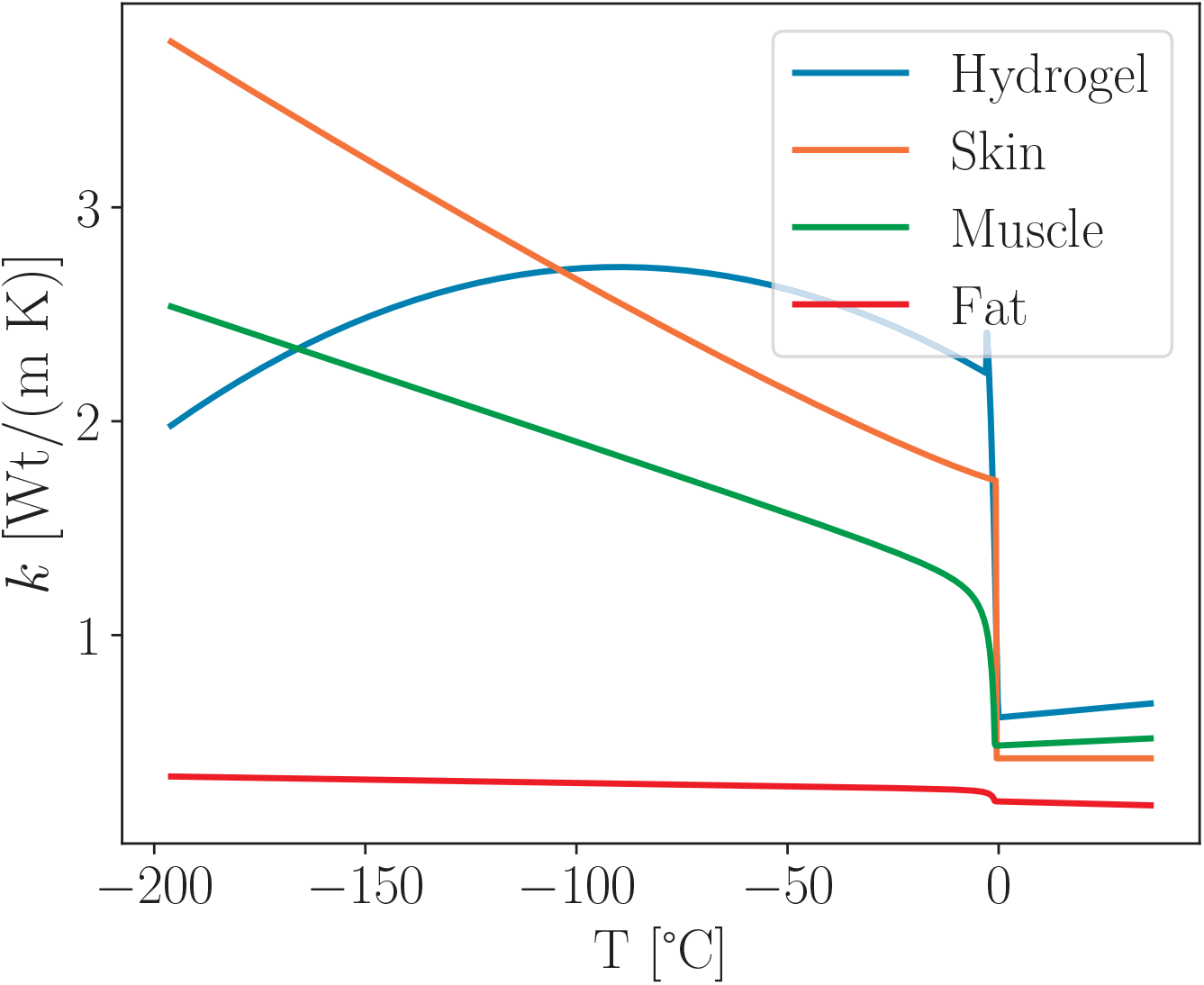
Temperature dependence of the thermal conductivity of the hydrogel and biological tissues, built based on Table 3 and Eq. (35).

In Table 4 we see different forms of the formulas for the thermal capacity of biological tissues. For the skin and under-skin fat, only the usual heat capacity is shown, and the heat capacity induced by latent phase change heat is defined by the same Eq. (38) as for hydrogel, where instead of *Lf*_FW_*w* corresponding latent heat from Table 1 is used. For muscles, latent heat is already included in the heat capacity, so Table 4 shows the effective thermal capacity for muscles.

**Table 4.**
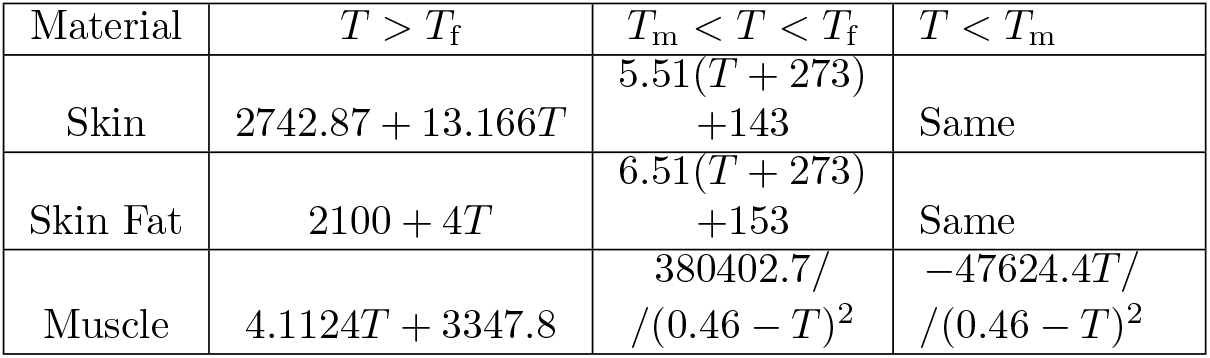
Thermal capacity for the biological tissues *C*_p_ [J/(kg K)] [28, 29].

| Material | $T > T_f$ | $T_m < T < T_f$ | $T < T_m$ |
| --- | --- | --- | --- |
| Skin | $2742.87 + 13.166T$ | $5.51(T + 273) + 143$ | Same |
| Skin Fat | $2100 + 4T$ | $6.51(T + 273) + 153$ | Same |
| Muscle | $4.1124T + 3347.8$ | $380402.7 / (0.46 - T)^2$ | $-47624.4T / (0.46 - T)^2$ |

In Fig. 12 we show the effective thermal capacity of the hydrogel and the biological tissues. There we find that the maximum value of the effective thermal capacity for the fat is similar to the maximum for other tissues despite it having around 2.5 times smaller latent heat. This corresponds to the fact that the range of the temperatures of the phase transition of the fat is smaller than for other materials, see Table 1. The total value of the latent heat is calculated as the integral of the latent heat over the phase change region for each tissue.

**Fig 12.**
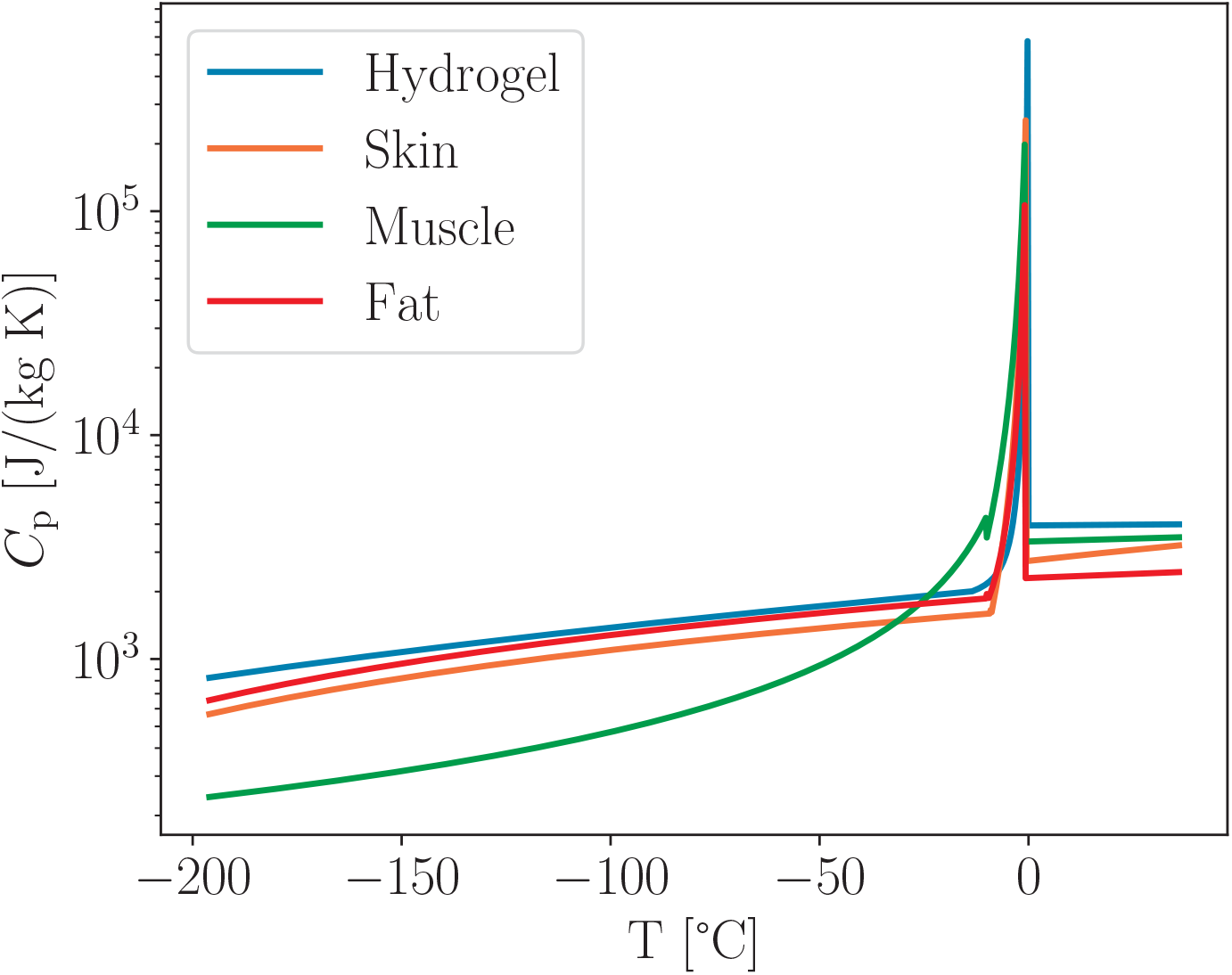
Temperature dependence of the effective thermal capacity for the biological tissues and hydrogel Eq. (38) (which includes latent phase change heat spread between *T*_m_ and *T*_f_ for each material)

## C Implementation of the finite difference method on GPU

To achieve high precision and fast calculating speed Graphics Processing Unit (GPU) can be very effective for large vectors like we use for the 2D cylindrical heat equation. GPU is very powerful for operations on multidimensional vectors with a huge number of elements because they are designed to calculate the color of each pixel on the computer screen, so for our calculations, we just use different algorithms to calculate color (temperature) in our simulation.

### C.1 Some features of solving heat equation on GPU

By utilizing the GPU power with optimizations, we achieve up to 20’000 times improvements in calculation speed in comparison to our calculation on the Intel I7 11900K CPU. It also allows us with powerful GPU like the Nvidia RTX 3080 for desktops to achieve real-time (sometimes even faster if the stability allows to increase timestep) calculation with high dimensional precision (in our case around 3.5 cm*/*512 = 0.06 mm per node), the main limitation usually is the stability of the FDM method [35]. This method becomes less stable with temperature-dependent coefficients like in our case. For our calculations, we use 512×512 nodes mesh, which means we have 262144 nodes. The typical time step we use is 10 microseconds, that means we have 100’000 time steps per second This means that to simulate dynamics during 1 second we update node temperature over 26 billion times. In some cases, our algorithm works stable and with high precision with up to 300 microseconds time step, which leads to a corresponding increase in calculating speed. The main limitation factor to increase the time step is the stability of the FDM method which can vary for the same dimensional steps if the thermodynamic parameters of the layers are significantly different.

To deal with these large arrays we minimized copy operations to use less memory. Specifically, we use the temperature nodes’ field with *N* + 2 dimensions to have an unused contour around the simulated area, so we can easily calculate neighbor nodes temperature and thermal conductivity from the same array just using shifted indexes. We use the thermal conductivity array to avoid additional calculations for the thermal conductivity in neighbor nodes, which updates for each node and then synchronizes.

We found that apart from the stability of the FDM, the main limitation is the memory transfer between the CPU and GPU, which is very slow compared to internal memory transfers within the GPU, so we draw the temperature field only every 10’000 time steps.

Avoiding redundant calculations is another way to enhance productivity, so the formulas for calculating on the GPU many times should be simplified as much as possible. For example, when we calculate the total heat capacity of a node *C* = *C*_p_*ρ* we can multiply these values and investigate the obtained graph, which could allow us to find a simpler formula for total heat capacity rather than calculate density *ρ* and specific heat capacity *C*_p_ and multiply them.

To perform the calculation for the heat equation with the FDM method we utilize the CuPy library for Python by writing CUDA rawkernel for the single time step update, this method allows us to avoid additional copying during the calculation CUDA kernel call.

To handle the border conditions we use two methods. First, we apply the zero conductivity to the nodes outside the calculated area, which with the mutual conductivity formula (29) gives us zero conductivity outside the calculation area. Second, we use the boolean masks for the thermal conductivity in certain directions, and this allows us to eliminate heat transfer from the sample to the cryoapplicator to keep the cryoapplicator temperature constant while including the heat transfer from the cryoapplicator to the sample.

Also, we use a mask for the materials that are used inside the kernel to identify the material and its properties.

### C.2 Kernel code sample

Here we present some useful parts of the CUDA kernel for the calculation of the single time-step. In the header of the RawKernel, we pass addresses of the arrays instead of copying arrays when calling the kernel by using the symbol ‘*’, also we use the modifier ‘const’ to be sure that we do not change values that are not supposed to be changed inside the kernel.

~~~
Calculate Frame Kernel_1=cp . RawKernel (r ‘ ‘ ‘
extern “C”{
_ _global_ _void Calculate Frame_Kernel1 (
float * T, float * K,
const float * rvals, const bool IsGel,
const float * dr,
const int * MaterialMask,
const bool * AlphDownMask,
const bool * AlphUpMask,
const bool * AlphRightMask,
const bool * AlphLeftMask)
{ \\ Kernel body
}
}
‘ ‘ ‘, ‘ Calculate Frame_Kernel 1 ‘)
~~~

Then we identify indexes *i, j* as well as global array index *idx* and indexes in the *N* + 2 dimensions to be able to use shifted coordinates. Also in this block, we unpack constants that we use later to calculate temperature-dependent thermodynamic parameters.

~~~
int id = blockIdx . x * blockDim . x
+ threadIdx . x ;
int jd = blockIdx . y * blockDim . y
+ threadIdx . y ;
int r_si z e=int (rvals[ 0 ]) ;
float dtdd=rvals [ 1 ] ;
inti=jd * r_size+id ;
int j = i%r_size;
int il =i / r_size+1;
int idx = il * (r_size+ 2) + j + 1 ;
const float rp=rvals [ j + 6 ] ;
const float rm=rvals [ j+6+r_size] ;
const float L_gelCor=rvals[ 2 ] ;
const float Tcor=rvals[ 3 ] ;
const float T_m_Gel=rvals[ 4 ] ;
const float T_f_Gel=rvals[ 5 ] ;
~~~

Then we take the temperature of the current node and the temperatures of the neighbor nodes and calculate their differences.

~~~
float Tij = T[ idx ] ;
const float T_up = T[ idx + r_size+ 2 ] ;
const float T_down = T[ idx − r_size− 2 ] ;
const float T_right= T[ idx + 1 ] ;
const float T_left = T[ idx − 1 ] ;
const float Tij_up = (T_up−Tij) ;
const float Tij_down = (T_down−Tij) ;
const float Tij_right= (T_right−Tij) ;
const float Tij left = (T_left −Tij) ;
~~~

Next, we calculate the thermal conductivity of the current node taking into account the material of the node, update it, and synchronize all GPU threads to ensure that the actual conductivity is used later.

~~~
float kij = computeGelCond (Tij, T_f_Gel, T_m_Gel,
L_gelCor, Tcor, IsGel, Material Mask [ idx ]) ;
K[ idx ]= k i j ;
_ _syncthreads () ;
~~~

After that, we take the conductivity of the neighboring nodes and calculate the heat capacity *C* = *C*_p_*ρ*.

~~~
float k_Up =K[ idx + r_size+ 2 ] ;
float k_Down =K[ idx − r_size− 2 ] ;
float k_Right =K[ idx + 1 ] ;
float k_Left =K[ idx − 1 ] ;
float Cij=Heat_Capacity (Tij, T_f_Gel, T_m_Gel,
L_gel Cor, Tcor, IsGel, Material Mask [ idx ]) ;
~~~

Then we calculate effective thermal diffusivity coefficients with all neighbors.

~~~
float GelAlphaUp =2*(kij *k_Up) /(kij+k_Up) /(Cij) ;
float GelAlphaDown =2*(kij *k_Down) /(kij+k_Down) /(Cij) ;
float Gel AlphaRight=
2 *(kij * k_Right) /(kij+k_Right) /(Cij) ;
float Gel Alpha Left =2*(kij * k_Left) /(kij+k_Left) /(Cij) ;
~~~

Next, we apply the half-step shifted radius for the radius-related nodes and apply masks to take into account border conditions for the cryoapplicator border.

~~~
float AlphRight = Gel AlphaRight* rp
*AlphRightMask [ i ] ;
float AlphLeft = Gel Alpha Left *rm ;
float AlphUp = GelAlphaUp* AlphUpMask [ i ] ;
float AlphDown = GelAlphaDown ;
~~~

Finally, we calculate the updated temperature field, where *dtdd* is the precalculated value for time step *dt* divided by depth step *dd, dtdd* = *dt/dd* to avoid redundant calculations.

~~~
Tij+= ((Tij_up *AlphUp +Tij_down *AlphDown) * dtdd
+ (Tij_right* AlphRight + Tij_left * AlphLeft)) ;
~~~

Then we synchronize all threads and update the temperature field array.

~~~
_ _syncthreads () ;
T[idx ]= Tij ;
~~~

As you can see from the code above, utilizing GPU power is not very difficult, but can give huge gains for the heat equation calculation, especially on the powerful GPU.

